# Reproducibility of R-fMRI Metrics on the Impact of Different Strategies for Multiple Comparison Correction and Sample Sizes

**DOI:** 10.1101/128645

**Authors:** Xiao Chen, Bin Lu, Chao-Gan Yan

## Abstract

Concerns regarding reproducibility of resting-state functional magnetic resonance imaging (R-fMRI) findings have been raised. Little is known about how to operationally define R-fMRI reproducibility and to what extent it is affected by multiple comparison correction strategies and sample size. We comprehensively assessed two aspects of reproducibility, test-retest reliability and replicability, on widely used R-fMRI metrics in both between-subject contrasts of sex differences and within-subject comparisons of eyes-open and eyes-closed (EOEC) conditions. We noted permutation test with Threshold-Free Cluster Enhancement (TFCE), a strict multiple comparison correction strategy, reached the best balance between family-wise error rate (under 5%) and test-retest reliability / replicability (e.g., 0.68 for test-retest reliability and 0.25 for replicability of amplitude of low-frequency fluctuations (ALFF) for between-subject sex differences, 0.49 for replicability of ALFF for within-subject EOEC differences). Although R-fMRI indices attained moderate reliabilities, they replicated poorly in distinct datasets (replicability < 0.3 for between-subject sex differences, < 0.5 for within-subject EOEC differences). By randomly drawing different sample sizes from a single site, we found reliability, sensitivity and positive predictive value (PPV) rose as sample size increased. Small sample sizes (e.g., < 80 (40 per group)) not only minimized power (sensitivity < 2%), but also decreased the likelihood that significant results reflect “true” effects (PPV < 0.26) in sex differences. Our findings have implications for how to select multiple comparison correction strategies and highlight the importance of sufficiently large sample sizes in R-fMRI studies to enhance reproducibility.

## 1. INTRODUCTION

The ability to replicate an entire experiment is essential to the scientific method (Open Science Collaboration, 2015). Much of the scientific enterprise, such as providing detailed descriptions of methods and peer-reviewing manuscripts before publication, is intended to optimize agreement of results when performed by different researchers. Such efforts are crucial because science cannot progress if results cannot be reproduced (Blackford, 2017). However, concerns regarding the reproducibility of biomedical and psychological research are increasingly being expressed (Open Science Collaboration, 2015; Ioannidis, 2005; Prinz, et al., 2011). This is particularly relevant to the field of resting-state functional magnetic resonance imaging (R-fMRI) (Carp, 2012a; Poldrack, et al., 2017), which has appeared to be a fruitful approach for basic, translational and clinical neuroscience (Biswal, et al., 1995; Fox and Raichle, 2007; Fox, et al., 2005). Beyond its reported sensitivity to developmental, aging and pathological processes (Hjelmervik, et al., 2014; Luo, et al., 2011; Tomasi and Volkow, 2012), R-fMRI is being increasingly adopted due to the relative ease of data collection and amenability to aggregation across studies and sites (Zuo, et al., 2014). These advantages are counterbalanced by high data dimensionality, relatively small sample size of most studies and the great amount of flexibility in data analysis, all of which threaten reproducibility.

Some aspects of R-fMRI reproducibility have been examined. Intra-class correlation (ICC), which models the ratio of between-subject variability to within-subject variability (Caceres, et al., 2009; Shrout and Fleiss, 1979), has been used to assess test-retest reliability, and moderate-to-high ICC has been reported for most R-fMRI metrics (Cao, et al., 2014; Shehzad, et al., 2009; Zuo and Xing, 2014; Zuo, et al., 2013; Zuo, et al., 2010a). However, ICC may be less informative, given the common practice in the field of reporting *P*- or *Z*-thresholded statistical maps (Kristo, et al., 2014). Different thresholding techniques vary in the sets and numbers of voxels, it is crucial to evaluate the test-rest reliability and replicability of the supra-threshold voxels. Thus, the first focus of the current study is to quantify the reliability replicability of R-fMRI metrics with thresholding. We compared differences of common metrics between males and females (between-subject) and between eyes-open (EO) and eyes-closed (EC) conditions (within-subject) and determined how well the significant clusters were reproduced on retests (test-retest reliability) or in totally different datasets/studies (replicability). Sex differences were chosen because sex is an objective category that can be readily investigated across large scale datasets. To wit, differences in brain function between men and women have been well documented in the R-fMRI literature (Allen, et al., 2011; et al., 2015; Bluhm, et al., 2008; Filippi, et al., 2013; Hjelmervik, et al., 2014; Kilpatrick, et al., 2015; Scheinost, et al., 2015; Tomasi and Volkow, 2012; Xu, et al., 2015). We chose to examine differences between EO and EC conditions to test whether our approach to within-subject designs. EO and EC differences have been reported to differ considerably in R-fMRI studies (Yan, et al., 2009; Zou, et al., 2009).

Test-rest reliability and replicability of the supra-threshold voxels are highly sensitive to the statistical threshold used to define significance. Such reproducibility was reported decreased as the significance threshold is enhanced (Duncan, et al., 2009). However, liberalizing the statistical threshold can dramatically increase the family-wise error rate (FWER), as recently demonstrated systematically for widely-used statistical methods (Eklund, et al., 2016). The trade-off between reproducibility and FWER requires a comprehensive investigation into different statistical approaches for multiple comparison correction to try to reach a balance. Accordingly, the impact of statistical method, especially multiple comparison correction strategies, on reproducibility is the second focus of the present study.

A third concern is the low statistical power of small samples, which are prevalent in the field of neuroscience. Carp reviewed over 200 fMRI studies published since 2007, and found the median sample size was 15 for one-group studies and 14.75 per group for two-group studies, resulting in unacceptable statistical power for most studies (Carp, 2012b). Another recent analysis (Poldrack, et al., 2017), reviewed 1131 sample sizes in neuroimaging studies over more than 20 years. Despite the steady increase in sample size (with median sample size up to 28.5 for single-group studies and 19 per group in multi-group studies), the median study in 2015 was only sufficiently powered to detect effects greater than 0.75. Button and colleagues calculated the statistical power of neuroscience studies with data extracted from meta-analyses. They found that the median statistical power of studies in the field of neuroscience was optimistically estimated to be between ∼8% and ∼31% (Button, et al., 2013). Moreover, the statistical findings of low power studies are unlikely to reflect true effects (i.e., they have low positive predictive value, PPV) (Button, et al., 2013; Ioannidis, 2005). Although these concerns have long been known, empirical evidence of how sample size influences reliability, as well as power and positive predictive value (PPV) of R-fMRI data are still scant. The attempt to quantify sensitivity and PPV has been hampered by the problem of how to define truly positive results. Here, we define findings that are reproducible in many datasets as the “gold standard”, which makes it possible to quantify sensitivity and PPV as a function of sample size.

To address the above issues, we systematically analyzed four independent datasets to quantify both the test-retest reliability and replicability of R-fMRI data and investigate how multiple comparison correction strategies impact them. We also considered how sample size might influence reliability as well as power and PPV. Five common R-fMRI metrics, namely, the amplitude of low frequency fluctuation (ALFF) and its fractional version (fALFF), regional homogeneity (ReHo), degree centrality (DC) and voxel-mirrored homotopic connectivity (VMHC) were employed to encompass possible sex and EOEC differences. We conclude by recommending a guideline based on this quantitative analysis to address the challenge of reproducibility in R-fMRI research.

## 2. MATERIALS AND METHODS

### 2.1. Participants and Imaging Protocols

We performed our analyses on four independent datasets. Three of them are publicly available via the International Neuroimaging Data-sharing Initiative (INDI, data available at http://fcon_1000.projects.nitrc.org): the Consortium for Reliability and Reproducibility (CORR) (Zuo, et al., 2014), the 1000 Functional Connectomes Project (FCP) (Biswal, et al., 2010) and Beijing EOEC1 (Liu, et al., 2013). The fourth dataset (Beijing EOEC2) was available through The R-fMRI Maps Project (http://rfmri.org/BeijingEOEC2_Raw), and was the basis of previous studies (Yan, et al., 2009; Zou, et al., 2009). The first two datasets were analyzed to evaluate test-retest reliability, replicability and the influence of sample size on between-subject sex differences (for details, see Tables S1-S2). The latter two datasets were employed to explore whether our approach generalizes to within-subject studies (EO and EC differences). In the former two datasets, participants were instructed to simply rest while awake in a scanner (mostly 3T, although three FCP sites used 1.5T scanners). In the latter two datasets, participants were instructed to open or to close their eyes while being scanned at 3T (8 minutes per session, EO and EC order counterbalanced across subjects). The R-fMRI data were acquired using an echo-planer imaging (EPI) sequence. A high-resolution T1-weighted anatomical image was also obtained for each participant for spatial normalization and localization. The corresponding institutional review boards of each collection center approved or provided waivers for the sharing of anonymized data, which were obtained with written informed consent from each participant.

The first dataset originally included 549 subjects who underwent 2 scanning sessions (mean time range = 205 ± 161 days) available at CORR. Of those, 420 subjects (age 21.45 ± 2.67, 208 females, henceforth the “CORR dataset”, see Table S1 for details) were selected after quality control with the following exclusion criteria. To avoid the confounds of development or aging, only young adults (age between 18 and 32) were included. Subjects were excluded if their functional scans showed excessive motion, indexed by mean frame-wise displacement (FD) (Jenkinson, et al., 2002) exceeding 0.2mm. Participants with poor T1 or functional images, low quality normalization or inadequate brain coverage were also excluded. The second dataset consisted of 716 young healthy subjects (age 22.34 ± 2.92, 420 females, henceforth the “FCP dataset”, see Table S2 for details) selected from FCP with the same inclusion criteria as the CORR dataset. The third dataset consisted of 48 healthy subjects (age 22.42 ± 2.24, 24 females, henceforth the “Beijing EOEC1 dataset”). The fourth dataset included 20 subjects (age 20.95 ± 1.82, 10 females, henceforth the “Beijing EOEC2 dataset”). The same inclusion criteria as the CORR and FCP datasets were applied, but no subject was excluded. For further information on the datasets including scanning protocols please refer to the CORR (http://fcon_1000.projects.nitrc.org/indi/CoRR/html/index.html), FCP (http://fcon_1000.projects.nitrc.org/index.html) and Beijing EOEC1 (http://fcon_1000.projects.nitrc.org/indi/retro/BeijingEOEC.html) websites. The Beijing EOEC2 dataset used the same scanning parameters as the Beijing EOEC1 dataset; the detailed protocol can be found in Yan et al. (2009).

### 2.2. Preprocessing

Unless otherwise stated, all preprocessing was performed using the Data Processing Assistant for Resting-State fMRI (DPARSF, Yan and Zang, 2010, http://rfmri.org/DPARSF), which is based on Statistical Parametric Mapping (SPM, http://www.fil.ion.ucl.ac.uk/spm) and the toolbox for Data Processing & Analysis of Brain Imaging (DPABI, Yan, et al., 2016, http://rfmri.org/DPABI). First, the initial 10 volumes were discarded, and slice-timing correction was performed with all volume slices corrected for different signal acquisition time by shifting the signal measured in each slice relative to the acquisition of the slice at the mid-point of each repetition time (TR). Then, the time series of images for each subject were realigned using a six-parameter (rigid body) linear transformation with a two-pass procedure (registered to the first image and then registered to the mean of the images after the first re-alignment). After realignment, individual T1-weighted MPRAGE images were co-registered to the mean functional image using a 6 degree-of-freedom linear transformation without re-sampling and then segmented into gray matter (GM), white matter (WM) and cerebrospinal fluid (CSF) (Ashburner and Friston, 2005). Finally, transformations from individual native space to MNI space were computed with the Diffeomorphic Anatomical Registration Through Exponentiated Lie algebra (DARTEL) tool (Ashburner, 2007).

### 2.3. Nuisance Regression

To minimize head motion confounds, we utilized the Friston 24-parameter model (Friston, et al., 1996) to regress out head motion effects. The Friston 24-parameter model (i.e., 6 head motion parameters, 6 head motion parameters one time point before, and the 12 corresponding squared items) was chosen based on prior work that higher-order models remove head motion effects better (Satterthwaite, et al., 2013; Yan, et al., 2013a). Additionally, mean FD was used to address the residual effects of motion in group analyses. Mean FD is derived from Jenkinson's relative root mean square (RMS) algorithm (Jenkinson, et al., 2002). As global signal regression (GSR) is still a controversial practice in the R-fMRI field, and given the recent advice that analyses with and without GSR be considered complementary (Murphy and Fox, 2016), we evaluated results both with and without GSR. Other sources of spurious variance (WM and CSF signals) were also removed from the data through linear regression to reduce respiratory and cardiac effects. Additionally, linear trends were included as a regressor to account for drifts in the blood oxygen level dependent (BOLD) signal. We performed temporal bandpass filtering (0.01-0.1Hz) on all time series except for ALFF and fALFF analyses. Of note, temporal bandpass filtering was performed after nuisance regression, thus would not reintroduce nuisance-related variation (Hallquist, et al., 2013).

### 2.4. A Broad Array of R-fMRI Metrics

Amplitude of Low Frequency Fluctuations (ALFF) (Zang, et al., 2007) and fractional ALFF (fALFF) (Zou, et al., 2008): ALFF is the mean of amplitudes within a specific frequency domain (here, 0.01-0.1Hz) from a fast Fourier transform of a voxel’s time course. fALFF is a normalized version of ALFF and represents the relative contribution of specific oscillations to the whole detectable frequency range.

Regional Homogeneity (ReHo) (Zang, et al., 2004): ReHo is a rank-based Kendall’s coefficient of concordance (KCC) that assesses the synchronization among a given voxel and its nearest neighbors’ (here, 26 voxels) time courses.

Degree Centrality (DC) (Buckner, et al., 2009; Zuo, et al., 2012): DC is the number or sum of weights of significant connections for a voxel. Here, we calculated the weighted sum of positive correlations by requiring each connection’s correlation coefficient to exceed a threshold of r > 0.25 (Buckner, et al., 2009).

Voxel-mirrored homotopic connectivity (VMHC, Anderson, et al., 2011; Zuo, et al., 2010b): VMHC corresponds to the functional connectivity between any pair of symmetric inter-hemispheric voxels - that is, the Pearson’s correlation coefficient between the time series of each voxel and that of its counterpart voxel at the same location in the opposite hemisphere. The resultant VMHC values were Fisher-Z transformed. For better correspondence between symmetric voxels, VMHC requires that individual functional data be further registered to a symmetric template and smoothed (4 mm FWHM). The group averaged symmetric template was created by first computing a mean normalized T1 image across participants, and then this image was averaged with its left–right mirrored version (Zuo, et al., 2010b).

Before entering into further analyses, all of the metric maps were Z-standardized (subtracting the mean value for the entire brain from each voxel, and dividing by the corresponding standard deviation) and then smoothed (4 mm FWHM), except for VMHC (which were smoothed and Fisher-Z transformed beforehand). To verify if our conclusions were affected by smoothing kernel, we have also re-analyzed our data with 8mm FWHM smoothing kernel.

### 2.5. Strategies to Correct for Multiple Comparisons

We first evaluated the FWER of 31 different statistical strategies (see Tables 1 and 2). Statistical maps were thresholded using eight versions of the one-tailed Gaussian random field theory (GRF) (Friston, et al., 1994; Nichols and Hayasaka, 2003) correction procedure, as implemented in DPABI (Yan, et al., 2016). These eight thresholding approaches used single-voxel thresholds (or cluster-defining thresholds) of *P* < 0.01 (*Z* > 2.33), *P* < 0.005 (*Z* > 2.58), *P* < 0.001 (*Z* > 3.09), or *P* < 0.0005 (*Z* > 3.29), and cluster size thresholds of *P* < 0.05, or *P* < 0.025. Given that GRF correction is only performed on one-tailed tests, we set *P* < 0.025 to perform two one-tailed tests, which is equivalent to two-tailed *P* < 0.05 after Bonferroni correction. Furthermore, we evaluated FWER of two versions of Monte Carlo simulation (simulated 1000 times) based corrections (Ledberg, et al., 1998), which is implemented in AFNI (AFNI 3dClustSim, https://afni.nimh.nih.gov/afni/doc/manual/3dclust.pdf) and DPABI (DPABI AlphaSim), separately. We note the bug reported in Eklund et al. (2016) had been fixed in the software versions used in the current study. Each version of Monte Carlo simulation based correction used the same eight thresholding approaches used for GRF. Statistical maps were also thresholded using seven kinds of permutation tests (PT), as implemented in PALM (Winkler, et al., 2016) and integrated into DPABI. For PALM approaches, two-tailed *P* < 0.05 (compared to 1000 permutations in FWER evaluation, and 5000 permutations for the remaining analyses) was set as the final threshold. For cluster-extent PT, voxel thresholds (cluster-defining thresholds) of two-tailed *P* < 0.02 (*Z* > 2.33), *P* < 0.01 (*Z* > 2.58), *P* < 0.002 (*Z* > 3.09) and *P* < 0.001 (*Z* > 3.29) were set. The threshold-free cluster enhancement (TFCE) (Smith and Nichols, 2009) and voxel-wise correction (VOX) with PT were also tested at two-tailed *P* < 0.05. Finally, false discovery rate (FDR) (Genovese, et al., 2002) correction was also examined.

**Table 1.** Family wise error rate and cluster size of ALFF (smoothness: 7.94×7.31×6.86) without GSR under corrections of Gaussian Random Field Theory, AFNI 3dClustSim and DPABI AlphaSim.

| Voxel Threshold<br>( One Tailed Twice ) | Cluster<br>Threshold | AFNI 3dClustSim |  | DPABI AlphaSim |  | Gaussian Random Field |  |
| --- | --- | --- | --- | --- | --- | --- | --- |
|  |  | Family Wise<br>Error Rate | Cluster Size | Family Wise<br>Error Rate | Cluster Size | Family Wise Error<br>Rate | Cluster Size |
| $P < 0.01$ ($Z > 2.33$) | $P < 0.05$ | 40.0% | 66.05±0.73 | 48.3% | 60.24±1.68 | 36.5% | 69.35±1.09 |
| $P < 0.005$ ($Z > 2.58$) | $P < 0.05$ | 27.6% | 43.59±0.42 | 34.9% | 39.45±1.13 | 24.5% | 46.70±0.75 |
| $P < 0.001$ ($Z > 3.09$) | $P < 0.05$ | 11.5% | 19.98±0.34 | 15.8% | 18.40±0.61 | 10.6% | 21.29±0.46 |
| $P < 0.0005$ ($Z > 3.29$) | $P < 0.05$ | 9.6% | 14.53±0.25 | 12.5% | 13.93±0.54 | 8.2% | 15.82±0.39 |
| $P < 0.01$ ($Z > 2.33$) | $P < 0.025$ | 30.8% | 74.50±1.14 | 39.0% | 67.72±2.36 | 27.7% | 78.96±1.24 |
| $P < 0.005$ ($Z > 2.58$) | $P < 0.025$ | 23.7% | 47.01±0.59 | 27.1% | 44.48±1.60 | 18.3% | 53.48±0.85 |
| $P < 0.001$ ($Z > 3.09$) | $P < 0.025$ | 8.6% | 22.63±0.25 | 10.6% | 21.00±0.87 | 6.8% | 24.94±0.41 |
| $P < 0.0005$ ($Z > 3.29$) | $P < 0.025$ | 5.8% | 17.33±0.22 | 7.9% | 16.03±0.71 | 5.1% | 18.51±0.50 |

**Table 2:** Family wise error rate under correction of 3 kinds of cluster-based correction with the strictest threshold, 6 versions of Permutation Test (PT) based correction as well as False Discovery Rate (FDR) correction. The smoothness in the second row is the estimated effective smoothness of the final metric maps feed to statistical analysis, and was different form the applied smoothness (4mm FWHM) in preprocessing. The effective smoothness was used in 3 versions of cluster-based correction (i.e., Gaussian Random Field theory correction, AFNI 3dClustSim and DPABI AlphaSim).

|  |  |  | Family Wise Error Rate |  |  |  |  |  |  |  |  |  |  |
| --- | --- | --- | --- | --- | --- | --- | --- | --- | --- | --- | --- | --- | --- |
|  | Voxel Threshold | Cluster Threshold | ALFF | fALFF | ReHo | DC | VMHC | ALFF with GSR | fALFF with GSR | ReHo with GSR | DC with GSR | VMHC with GSR | ALFF (8mm smoothed) |
|  | Smoothness (mm, x×y×z) |  | 7.94×7.31 | 7.34×7.42 | 9.36×8.72 | 7.86×7.97 | 6.31×6.87 | 7.99×7.31 | 7.32×7.41 | 9.24×8.56 | 8.06×8.16 | 6.11×6.61 | 11.88×11.53 |
|  |  |  | ×6.86 | ×7.20 | ×8.39 | ×7.81 | ×6.61 | ×6.84 | ×7.19 | ×8.18 | ×8.09 | ×6.37 | ×11.68 |
| AFNI 3dClustSim |  |  | 5.8% | 6.1% | 7.3% | 8.5% | 6.0% | 5.3% | 6.6% | 6.9% | 6.8% | 6.4% | 5.5% |
| (One Tailed) |  |  |  |  |  |  |  |  |  |  |  |  |  |
| DPABI AlphaSim | $P < 0.0005$ ($Z > 3.29$) | $P < 0.025$ | 7.9% | 8.3% | 8.5% | 10.2% | 9.0% | 7.8% | 7.7% | 7.8% | 8.3% | 9.6% | 6.9% |
| (One Tailed) |  |  |  |  |  |  |  |  |  |  |  |  |  |
| Gaussian Random |  |  | 5.1% | 5.5% | 4.9% | 7.4% | 5.2% | 4.8% | 5.9% | 5.3% | 5.1% | 6.4% | 4.4% |
| Field (One Tailed) |  |  |  |  |  |  |  |  |  |  |  |  |  |
| PT Cluster Extent | $P < 0.02$ ($Z > 2.33$) | $P < 0.05$ | 5.8% | 3.6% | 5.8% | 4.6% | 5.2% | 4.8% | 3.9% | 5.5% | 5.2% | 4.3% | 5.3% |
| Correction (Two |  |  |  |  |  |  |  |  |  |  |  |  |  |
| Tailed) | $P < 0.01$ ($Z > 2.58$) | $P < 0.05$ | 5.4% | 4.0% | 5.7% | 4.6% | 5.5% | 5.3% | 3.8% | 5.3% | 5.0% | 4.5% | 5.4% |
| | $P < 0.002$ ($Z > 3.09$) | $P < 0.05$ | 4.5% | 4.1% | 5.3% | 4.8% | 4.2% | 4.5% | 5.0% | 5.1% | 4.7% | 4.3% | 4.4% |
| | $P < 0.001$ ($Z > 3.29$) | $P < 0.05$ | 4.8% | 4.5% | 4.5% | 4.9% | 3.4% | 4.3% | 4.8% | 5.4% | 4.2% | 3.9% | 4.1% |
| PT Threshold-Free Cluster Enhancement (TFCE) |  |  | 4.6% | 3.9% | 5.7% | 5.0% | 4.3% | 5.3% | 4.2% | 5.5% | 4.7% | 4.8% | 4.6% |
| PT Voxel-Wise Correction (VOX) |  |  | 4.9% | 4.9% | 5.7% | 3.9% | 4.7% | 6.0% | 4.5% | 5.6% | 4.0% | 4.6% | 3.9% |
| FDR Correction |  |  | 3.1% | 3.4% | 4.4% | 2.4% | 3.9% | 4.1% | 2.8% | 3.6% | 2.4% | 3.5% | 1.6% |

### 2.6. Evaluating FWER of Different Strategies to Correct for Multiple Comparisons

To calculate the FWERs of different approaches for multiple comparison corrections, we performed permutation tests (1000 permutations in this study). For this permutation test, we first selected 106 female young subjects from the Beijing site within the FCP dataset to maximize sample homogeneity. Then, 40 subjects were randomly picked from the set of 106 subjects and randomly assigned to two equal groups (20 per group). Because assignment was fully random, no significant results should have emerged when these two groups’ R-fMRI metrics were compared. Detection of a significant difference after multiple comparison correction indicated a family wise error had occurred. Thus, FWER was calculated as the proportion of such false positives in all comparisons within the permutation test.

### 2.7. Assessing Test-Retest Reliability and Replicability of Different Datasets with Regard to Between-Subject Sex Differences and Within-Subject EOEC Differences

We first assessed the test-retest reliability and replicability of sex differences with CORR and FCP datasets. For each of the first two datasets, we employed a general linear model to examine the sex differences in R-fMRI measures while taking the confounding effects of age, head motion (mean FD) and site into account. Sex effect was estimated by the *t* value of the regressor corresponding to sex. Then the group difference map was corrected using different multiple comparison correction approaches described above to obtain statistically significant clusters.

The Dice coefficient was used to evaluate test-retest reliability as calculated by the following equation:

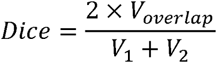

Where V_1_ and V_2_ represents the number of supra-threshold voxels in test 1 and test 2 of the CORR dataset, and V_overlap_ stands for the number of supra-threshold voxels in both tests.

To calculate replicability, we selected the voxels which were significant in both sessions in the CORR dataset, and then calculated how they overlapped with the significant voxels in the FCP dataset. We used the same Dice formula, with V_1_ representing the number of voxels significant in both sessions in the CORR dataset, V_2_ representing the number of voxels significant in the FCP dataset, and V_overlap_ standing for the number of voxels that were significant in both sessions in the CORR dataset as well as significant in the FCP dataset.

For each multiple comparison correction strategy, we calculated test-retest reliability and replicability. To figure out which multiple comparison correction strategy yielded the best test-retest reliability and replicability, a non-parametric one-way repeated measures ANOVA (Friedman’s test) on 5 metrics by 2 preprocessing strategies (with and without GSR) was conducted, and followed by post-hoc analyses corrected by Tukey's honest significant difference criterion.

Finally, we defined the voxels that were significant in both CORR sessions and in the FCP dataset as the “gold standard” for further evaluation (see section below). We believe these consistently significant voxels reflect true differences between males and females based on their high test-retest reliability and replicability in two large sample datasets.

To see whether our findings on between-subject sex differences generalized to within-subject EOEC differences, we further evaluated the replicability of the EOEC differences across two Beijing EOEC datasets. For each dataset, paired-t tests between EC and EO conditions were performed to examine EOEC differences in R-fMRI measures, while taking the confounding effect of head motion (mean FD) into account. Of note, between-subject factors (e.g., age and sex) did not need to be covaried in this within-subject design. Then, the EOEC difference map was corrected using the different previously described multiple comparison correction approaches to obtain statistically significant clusters. Similar to the sex difference analyses, the Dice coefficient was employed to calculate the replicability of EOEC differences between two Beijing EOEC datasets. Then, a non-parametric one-way repeated measures ANOVA (Friedman’s test) on 5 metrics by 2 preprocessing strategies (with and without GSR) and post-hoc analyses corrected by Tukey's honest significant difference criterion were conducted to evaluate all multiple comparison correction strategies with regard to replicability of EOEC differences.

### 2.8. Influences of Sample Size on Test-retest Reliability, Sensitivity and Positive Predictive Value

To estimate the influence of sample size on test-retest reliability, we tested the Dice coefficient of two tests (test/retest) as a function of sample size (k∈{30,40,50,60,70,80,90,100,120,140,160,180,200}). First, we randomized 100 times the order of female participants (and separately the order of male participants) from a single site (the “SWU 4” site in the CORR dataset, which has two sessions of 116 males and 105 females, for details see Table S1). Second, for each randomization of each k, we selected the first k/2 female participants and the first k/2 male participants. We then performed two-sample t-tests on the ALFF maps (without GSR) between males and females (with age and head motion as covariates) and then applied permutation test with TFCE (which performs better, see Results) to threshold the results. Finally, we calculated the Dice coefficient between the thresholded maps (binarized) of the first test and the retest, for each of the 100 randomizations and each k.

We also evaluated the sensitivity and PPV of the voxels which were significant in both tests of each randomization and each k, based on the “gold standard” defined in the prior section. The sensitivity of a study measures the proportion of positives that were correctly identified as such (Altman and Bland, 1994), while PPV is the probability that a positive finding reflects a true effect (Ioannidis, 2005). A recent analysis (Button, et al., 2013) demonstrated that studies with small sample size not only reduce the chance of detecting a true effect, but also reduce the probability that significant findings reflect a true effect. To determine this effect of sample size, the sensitivity and PPV were calculated:

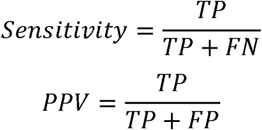

where TP is equal to the number of “true positive voxels”, which were statistically significant and reflected the true effect. As the true effect is difficult to define, we used the voxels that were significant in both CORR sessions and in the FCP dataset (the “gold standard” voxels defined above) after PT correction with TFCE. FN represents the number of “false negative voxels” that were statistically insignificant but reflected a true effect. And FP stands for the number of the false positive voxels that were statistically significant but did not reflect a true effect.

Of note, the key source codes for the analyses of the current study have been released at https://github.com/Chaogan-Yan/PaperScripts/tree/master/Chen_2017_HBM, thus readers can check, replicate or make use of our scripts in their future studies. In addition, maps of all the R-fMRI metrics of the four datasets used in the current study have been openly shared through the R-fMRI Maps Project (http://rfmri.org/maps), thus readers can easily replicate the current results based on these shared maps.

## 3. RESULTS

### 3.1. FWER of Different Multiple Comparison Correction Strategies

To evaluate the test-retest reliability and replicability of R-fMRI metrics, an appropriate statistical threshold and multiple comparison correction strategy must be defined in advance. The appropriate multiple comparison correction strategy must control the false positive rate at an acceptable level. Here, we evaluated 31 different multiple comparison correction strategies with 5 different R-fMRI metrics by 2 different preprocessing strategies (with and without GSR) in 106 female young adults (selected from the Beijing site of the FCP dataset). Based on the group differences of two randomly assigned groups (20 subjects per group, permuted 1000 times), we calculated FWER for each multiple comparison correction strategy. Table 1 presents FWERs and cluster sizes of GRF and Monte Carlo Simulation based correction strategies on ALFF without GSR. FWERs of other metrics with and without GSR can be found in supplementary materials (Tables S3-S11). For FWERs under GRF and Monte Carlo Simulation based corrections, the liberal voxel *P* thresholds (cluster-defining threshold) (*P* < 0.01 (*Z* > 2.33) and *P* < 0.005 (*Z* > 2.58)) far exceeded nominal 5% level (Table 1, Figure 1 & Tables S3-S11). Furthermore, as most researchers are interested in two-tailed effects (e.g., both patients > controls and patients < controls), if they perform one-tailed thresholding twice (i.e., each tail *P* < 0.05), then the final FWER is higher than the nominal 5% level. Only if the researcher corrects the two tests of each tail (e.g., Bonferroni correction, each tail controlled at *P* < 0.025), can the FWER reach the nominal 5% level. For example, GRF was almost valid under the strictest threshold (voxel-wise *P* < 0.0005 and cluster-wise *P* < 0.025, each tail): FWER of all metrics under 6.35% (the upper limit of approximate theoretical 95% confidence interval of nominal 5%) except for DC (Table 2). However, for Monte Carlo Simulation based corrections, the cluster size for thresholding is looser (smaller) than GRF (Tables 1, S3-S11), thus the FWER is higher than GRF: there were much more FWERs exceeded 6.35% even under the strictest threshold, especially for DPABI AlphaSim (Table 2). These findings were replicated when we re-analyzed the data with 8mm FWMH smoothing kernel in preprocessing (Figure S1, Table 2 last column and Table S12).

**Figure 1.**
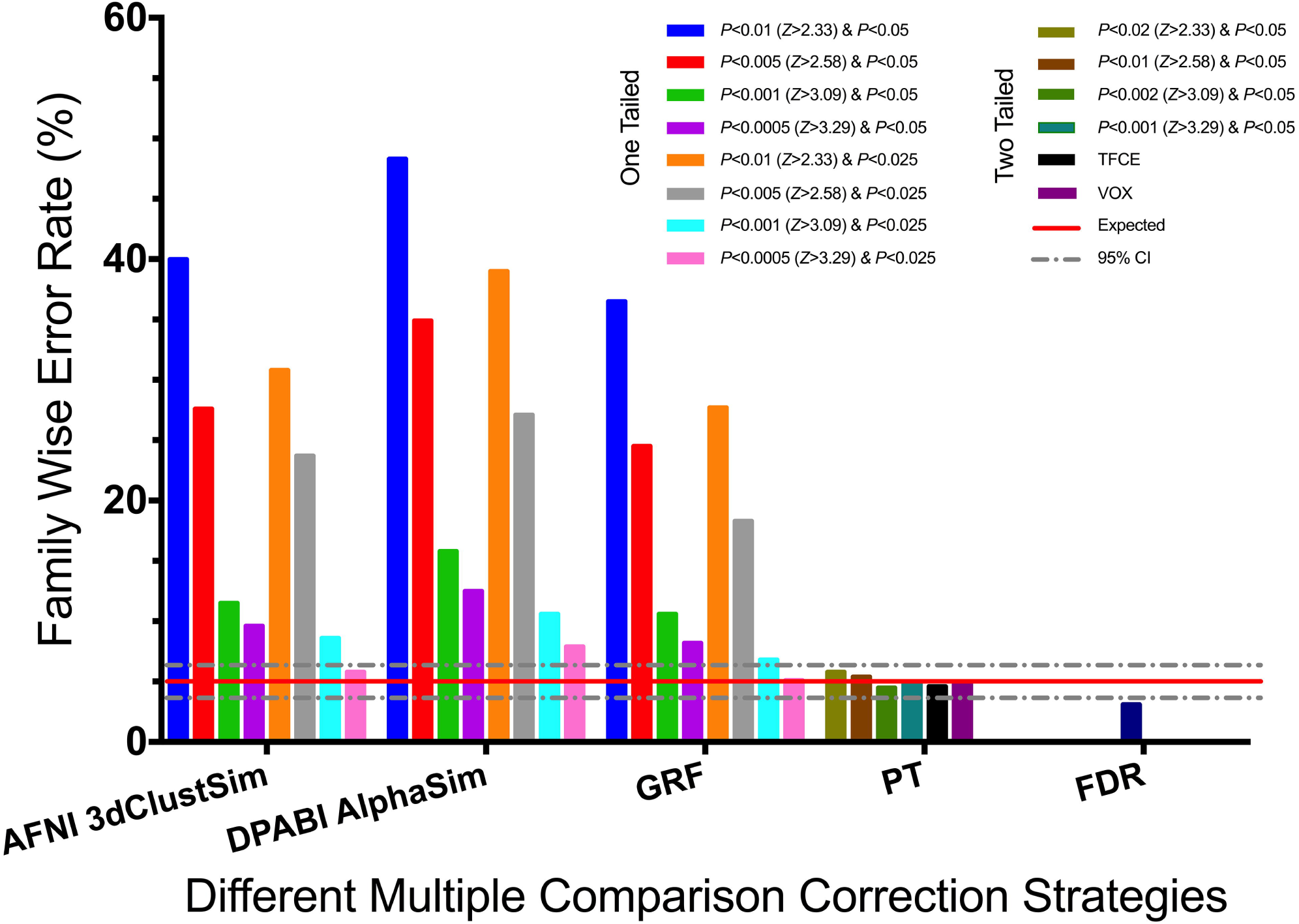
Family wise error rates of ALFF (without GSR) under 31 kinds of different multiple comparison correction strategies. AFNI 3dClustSim and DPABI AlphaSim are two versions of Monte Carlo simulation based correction implemented in AFNI and DPABI, separately. GRF, PT and FDR are Gaussian Random Field correction, Permutation Test and False Discovery Rate correction implemented in DPABI, separately. TFCE stands for Threshold-Free Cluster Enhancement and VOX stands for Voxel-Wise Correction. Both of them are correction approaches accompanied with PT. The red solid line shows the nominal 5% positive false positive rate, and the gray dashed line shows its approximate theoretical 95% confidence interval, 3.65%∼6.35%.

### 3.2. Test-retest Reliabilities of R-fMRI Metrics under Different Multiple Comparison Correction Strategies with Regard to Between-Subject Sex Differences

After evaluating the FWER, we systematically evaluated the test-retest reliabilities of five R-fMRI metrics under 31 different multiple comparison correction strategies on the CORR dataset (Tables 3 and S13). On average, test-retest reliability reached 0.50 (SD: 0.13, Range: 0.11 ∼ 0.75) among different R-fMRI metrics. ALFF, fALFF and ReHo had relatively high test-retest reliabilities: ALFF: 0.66 ± 0.01, fALFF: 0.61 ± 0.09, ReHo: 0.54 ± 0.05. In contrast, DC and VMHC had lower test-retest reliabilities: DC: 0.39 ± 0.05, VMHC: 0.43 ± 0.06. Interestingly, we found GSR decreased the test-retest reliability of R-fMRI metrics. For example, the test-retest reliability of DC decreased from 0.48 to 0.31 under correction of PT with TFCE when computed with GSR.

**Table 3.** Test-retest reliability of sex differences for all R-fMRI metrics with and without Global Signal Regression (GSR) under correction of 3 kinds of cluster-based correction with the strictest threshold, 6 kinds of Permutation Test (PT) based correction and False Discovery Rate (FDR) correction, calculated between the first and second sessions in the CORR dataset. For test-retest reliability for all the 31 kinds of multiple comparison correction strategies, please see Table S13.

|  |  |  | Test-retest Reliability (Dice Coefficient) |  |  |  |  |  |  |  |  |  |
| --- | --- | --- | --- | --- | --- | --- | --- | --- | --- | --- | --- | --- |
|  | Voxel Threshold | Cluster Threshold | ALFF | fALFF | ReHo | DC | VMHC | ALFF with GSR | fALFF with GSR | ReHo with GSR | DC with GSR | VMHC with GSR |
| AFNI 3dClustSim |  |  |  |  |  |  |  |  |  |  |  |  |
| (One Tailed) |  |  | 0.65 | 0.51 | 0.50 | 0.34 | 0.39 | 0.64 | 0.48 | 0.44 | 0.28 | 0.24 |
| DPABI AlphaSim | $P < 0.0005$ ($Z > 3.29$) | $P < 0.025$ | | | | | | | | | | |
| (One Tailed) |  |  | 0.65 | 0.51 | 0.49 | 0.34 | 0.39 | 0.64 | 0.48 | 0.45 | 0.27 | 0.27 |
| GRF (One Tailed) |  |  | 0.64 | 0.51 | 0.50 | 0.35 | 0.39 | 0.65 | 0.48 | 0.43 | 0.28 | 0.24 |
| PT Cluster Extent | $P < 0.02$ ($Z > 2.33$) | $P < 0.05$ | 0.65 | 0.70 | 0.56 | 0.45 | 0.40 | 0.62 | 0.68 | 0.45 | 0.30 | 0.40 |
| Correction (Two Tailed) | $P < 0.01$ ($Z > 2.58$) | $P < 0.05$ | 0.67 | 0.66 | 0.52 | 0.32 | 0.33 | 0.60 | 0.63 | 0.46 | 0.27 | 0.32 |
| | $P < 0.002$ ($Z > 3.09$) | $P < 0.05$ | 0.63 | 0.55 | 0.51 | 0.36 | 0.38 | 0.63 | 0.52 | 0.47 | 0.23 | 0.32 |
| | $P < 0.001$ ($Z > 3.29$) | $P < 0.05$ | 0.64 | 0.51 | 0.48 | 0.37 | 0.38 | 0.64 | 0.48 | 0.44 | 0.28 | 0.26 |
| PT Threshold-Free Cluster Enhancement (TFCE) |  |  | 0.68 | 0.75 | 0.54 | 0.48 | 0.44 | 0.66 | 0.74 | 0.44 | 0.31 | 0.42 |
| PT Voxel-Wise Correction (VOX) |  |  | 0.66 | 0.34 | 0.48 | 0.37 | 0.22 | 0.65 | 0.31 | 0.38 | 0.11 | 0.14 |
| FDR Correction |  |  | 0.64 | 0.67 | 0.54 | 0.39 | 0.37 | 0.63 | 0.64 | 0.47 | 0.23 | 0.29 |

Among those multiple comparison correction strategies that can control FWER under nominal 5%, we would like to identify which one could achieve the best test-retest reliability. We performed a Friedman test on 5 metrics by 2 preprocessing strategies (with and without GSR) among 10 strategies of multiple comparison correction: 3 kinds of cluster-based correction (i.e., GRF, AFNI 3dClustSim and DPABI AlphaSim, the latter two were added for comparison although they cannot always control FWER under nominal 5%) with the strictest threshold (voxel-wise *P* < 0.0005 and cluster-wise *P* < 0.025, each tail), 6 kinds of PT and FDR (Figure 2A). The 10 different multiple comparison correction strategies differed significantly (Friedman’s chi-square = 35.04, df = 9, *P* < 0.0001). Further post-hoc analysis revealed that PT with TFCE achieved the best test-retest reliability. PT with TFCE had significantly higher test-retest reliability than 3 kinds of cluster-based correction (i.e., GRF, AFNI 3dClustSim and DPABI AlphaSim) with the strictest threshold (voxel-wise *P* < 0.0005 and cluster-wise *P* < 0.025, each tail), PT (voxel-wise threshold of *P* < 0.002 (Z > 3.09) and *P* < 0.001 (Z > 3.29) with cluster-wise thresholds of *P* < 0.05 (two tailed)) and PT with voxel-wise correction (VOX) in the post-hoc analysis (*P* < 0.05, multiple comparison corrected by Tukey's honest significant difference criterion) (Figure 2A).

**Figure 2.**
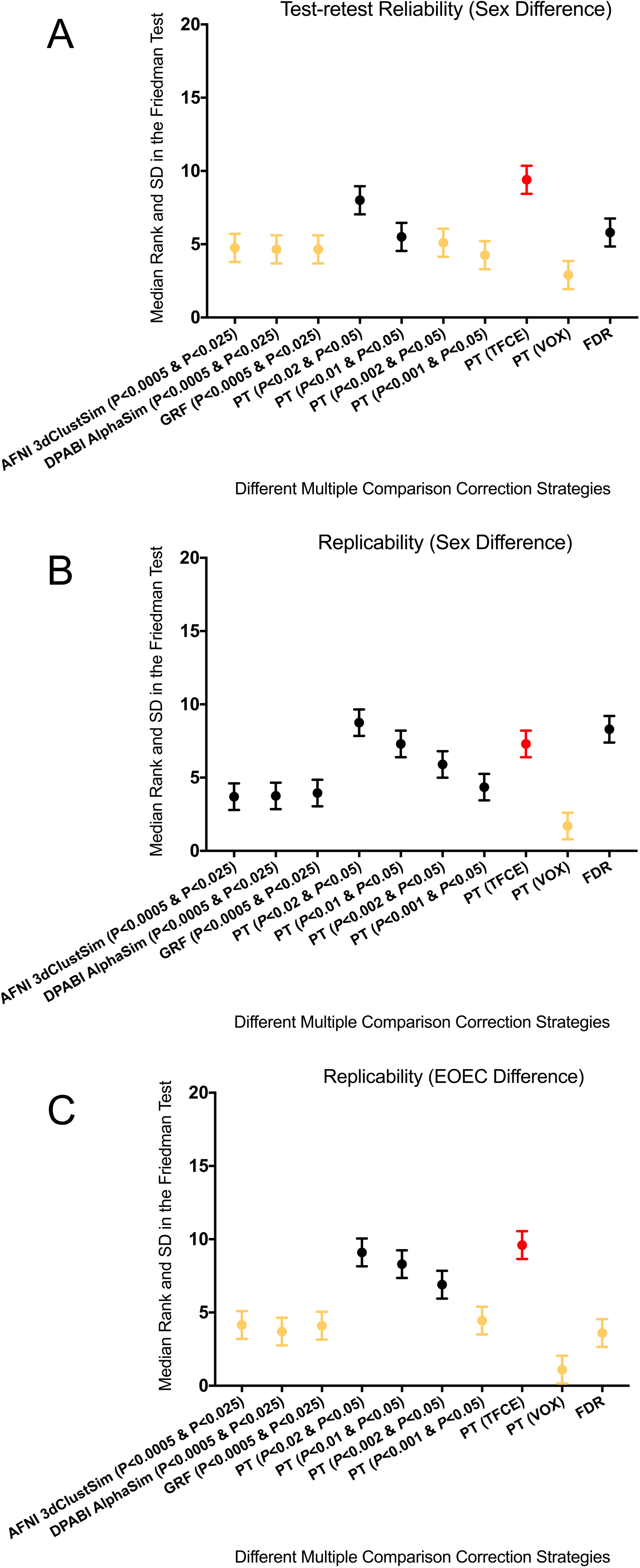
Results of the Friedman Test of both test-retest reliabilities and replicabilities regarding between-subject sex differences and within-subject eyes-open eyes-closed (EOEC) differences on 5 metrics by 2 preprocessing strategies (with and without GSR) among 3 kinds of cluster-based correction with the strictest threshold, 6 kinds of Permutation Test (PT) based correction and False Discovery Rate (FDR) correction (A: test-retest reliability regarding between-subject sex differences B: replicability regarding between-subject sex differences C: replicability regarding within-subject EOEC differences). Larger median rank numbers represent the better reproducibility compared with other statistical threshold approaches. PT with TFCE is outlined with red, and those are significantly different from PT with TFCE in reproducibility are outlined with yellow (multiple comparison corrected by Tukey's honest significant difference criterion). GRF, PT and FDR stand for Gaussian Random Field correction, Permutation Test and False Discovery Rate correction, separately. All versions of cluster-based corrections are one-tailed *P* values while all versions of PT are two tailed *P* values.

In addition, we found test-retest reliabilities under cluster-based correction (i.e., GRF, AFNI 3dClustSim and DPABI AlphaSim) with looser thresholds (e.g., voxel-wise *P* < 0.01 with cluster-wise *P* < 0.05, each tail) was higher than those with stricter thresholds (e.g., voxel-wise *P* < 0.0005 and cluster-wise *P* < 0.025, each tail) (Table S13). However, even at the cost of high FWER, cluster-based correction with loose thresholds did not show significantly higher test-retest reliability than PT with TFCE. Thus we conclude PT with TFCE was best able to balance FWER and test-retest reliability.

### 3.3. Replicability of R-fMRI Metrics under Different Multiple Comparison Correction Strategies with Regard to Between-Subject Sex Differences

To calculate replicability, we selected the voxels that were significant in both CORR sessions, and then calculated their overlap with the significant voxels in the FCP dataset (Tables 4 and S14). Generally, replicability was lower than test-retest reliability, achieving a mean of 0.11 (SD: 0.06, Range: 0.00 ∼ 0.28). Under the multiple comparison correction of PT with TFCE, ALFF (without GSR) reached a replicability of 0.25. None of the measures reached replicability higher than 0.3. This means that even voxels that could be reliably detected in two different sessions in the same subjects were difficult to replicate in a totally different dataset. This might be due to the many different factors between the two different datasets, for example, variation in ethnicity, sequence type, coil type, scanning parameters, participant instructions, head-motion restraint techniques, etc.

**Table 4.**
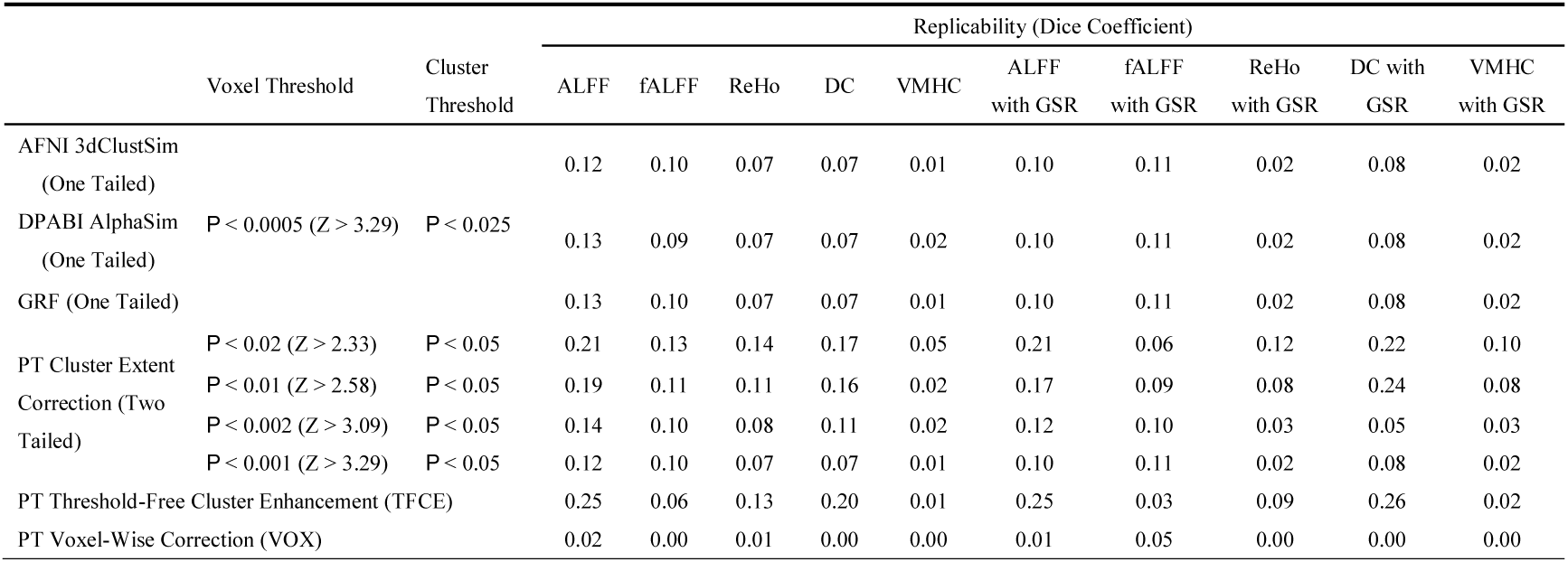
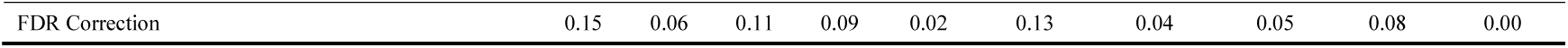
Replicability of sex differences for all R-fMRI metrics with and without Global Signal Regression (GSR) under correction of 3 kinds of cluster-based correction with the strictest threshold, 6 kinds of Permutation Test (PT) based correction and False Discovery Rate (FDR) correction, calculated using significant results in both sessions in the CORR dataset and those significant in the FCP dataset. For replicability for all the 31 kinds of multiple comparison correction strategies, please see Table S14.

|  | Voxel Threshold | Cluster Threshold | Replicability (Dice Coefficient) |  |  |  |  |  |  |  |  |  |
| --- | --- | --- | --- | --- | --- | --- | --- | --- | --- | --- | --- | --- |
|  |  |  | ALFF | fALFF | ReHo | DC | VMHC | ALFF with GSR | fALFF with GSR | ReHo with GSR | DC with GSR | VMHC with GSR |
| AFNI 3dClustSim (One Tailed) |  |  | 0.12 | 0.10 | 0.07 | 0.07 | 0.01 | 0.10 | 0.11 | 0.02 | 0.08 | 0.02 |
| DPABI AlphaSim (One Tailed) | $P < 0.0005$ ($Z > 3.29$) | $P < 0.025$ | 0.13 | 0.09 | 0.07 | 0.07 | 0.02 | 0.10 | 0.11 | 0.02 | 0.08 | 0.02 |
| GRF (One Tailed) |  |  | 0.13 | 0.10 | 0.07 | 0.07 | 0.01 | 0.10 | 0.11 | 0.02 | 0.08 | 0.02 |
| PT Cluster Extent | $P < 0.02$ ($Z > 2.33$) | $P < 0.05$ | 0.21 | 0.13 | 0.14 | 0.17 | 0.05 | 0.21 | 0.06 | 0.12 | 0.22 | 0.10 |
| Correction (Two Tailed) | $P < 0.01$ ($Z > 2.58$) | $P < 0.05$ | 0.19 | 0.11 | 0.11 | 0.16 | 0.02 | 0.17 | 0.09 | 0.08 | 0.24 | 0.08 |
| | $P < 0.002$ ($Z > 3.09$) | $P < 0.05$ | 0.14 | 0.10 | 0.08 | 0.11 | 0.02 | 0.12 | 0.10 | 0.03 | 0.05 | 0.03 |
| | $P < 0.001$ ($Z > 3.29$) | $P < 0.05$ | 0.12 | 0.10 | 0.07 | 0.07 | 0.01 | 0.10 | 0.11 | 0.02 | 0.08 | 0.02 |
| PT Threshold-Free Cluster Enhancement (TFCE) |  |  | 0.25 | 0.06 | 0.13 | 0.20 | 0.01 | 0.25 | 0.03 | 0.09 | 0.26 | 0.02 |
| PT Voxel-Wise Correction (VOX) |  |  | 0.02 | 0.00 | 0.01 | 0.00 | 0.00 | 0.01 | 0.05 | 0.00 | 0.00 | 0.00 |

|  |  |  |  |  |  |  |  |  |  |  |
| --- | --- | --- | --- | --- | --- | --- | --- | --- | --- | --- |
| FDR Correction | 0.15 | 0.06 | 0.11 | 0.09 | 0.02 | 0.13 | 0.04 | 0.05 | 0.08 | 0.00 |

A Friedman’s test was conducted to compare replicability under the abovementioned 10 different multiple comparison correction strategies (Figure 2B). The 10 different multiple comparison correction strategies differed significantly (Friedman’s chi-square = 45.73, df = 9, *P* < 10^-6^). PT with TFCE had significantly higher replicability than PT with VOX in post-hoc analysis (*P* < 0.05, multiple comparison corrected by Tukey's honest significant difference criterion) (Figure 2B). Again, we found that, even at the cost of high FWER, cluster-based correction with liberal thresholds (voxel-wise *P* < 0.01 and cluster-wise *P* < 0.05, each tail) did not show significantly higher replicability than PT with TFCE (Table S14).

### 3.4. Core Brain Regions with Reliable and Replicable Sex Differences

Sections 3.1 ∼ 3.3 showed that PT with TFCE yielded moderate test-retest reliability and replicability while maintaining FWER under the nominal 5% level, thus outperforming the alternative multiple comparison correction strategies. This allowed us to determine the core brain regions which differ by sex in R-fMRI metrics by identifying voxels that were replicated across both sessions of the CORR dataset and the FCP dataset when applying PT with TFCE correction. As shown in Figure 3, significant differences between males and females were reproducibly observed for all R-fMRI metrics. Brain regions with sex differences varied across R-fMRI metrics, although they converged at the posterior cingulate cortex (PCC). PCC demonstrated lower spontaneous activity in males compared with females in all the metrics except for DC (i.e., ALFF, fALFF, ReHo and VMHC). The voxels with replicable sex differences were considered the “gold standard” in Section 3.6 to calculate sensitivity and PPV with different sample sizes.

**Figure 3.**
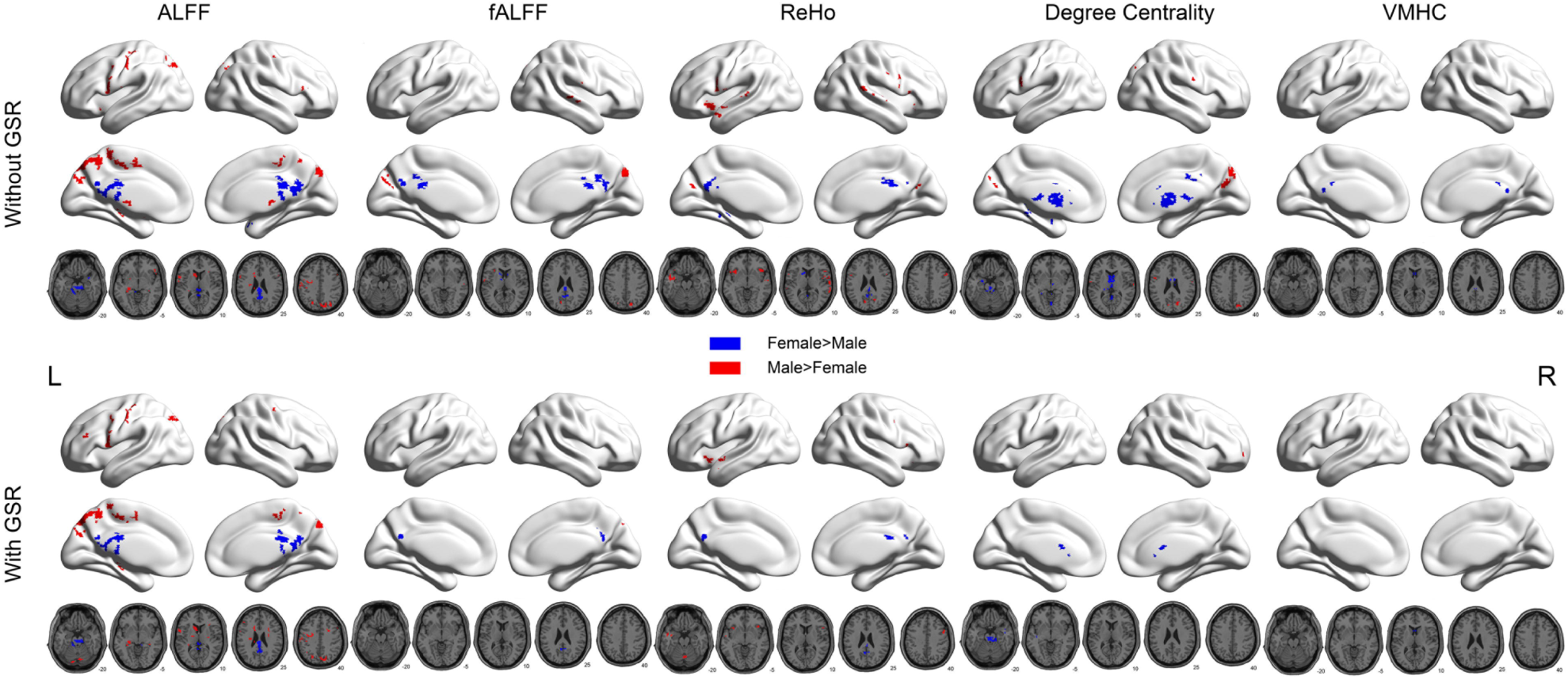
Sex differences those are significant in both sessions in the CORR dataset as well as significant in the FCP dataset (“gold standard”), under the correction of Permutation Test (PT) with Threshold-Free Cluster Enhancement (TFCE).

### 3.5. Replicability of R-fMRI Metrics under Different Multiple Comparison Correction Strategies with Regard to Within-Subject EOEC Differences

To verify whether our approach generalizes to within-subject design studies, we further calculated replicability of significant voxels from two EOEC datasets (Beijing EOEC1 and EOEC2 datasets). Again, we used the Dice coefficient to evaluate replicability (see Tables 5 and S15). Although replicabilities of within-subject EOEC differences were higher than between-subject sex differences (Mean: 0.23, SD: 0.13, Range: 0.00 ∼ 0.50), overall replicability still did not reach adequate levels. Similarly, replicabilities of ALFF (0.30 ± 0.14), fALFF (0.19 ± 0.09) and ReHo (0.35 ± 0.10) were higher than those of DC (0.12 ± 0.09) and VMHC (0.16 ± 0.07). We then conducted a Friedman’s test to compare replicability under the abovementioned 10 different multiple comparison correction strategies. The different multiple comparison correction strategies differed significantly (Friedman’s chi-square = 78.61, df = 9, P < 10^-12^). Again, post-hoc analysis revealed that PT with TFCE had the best replicability (Figure 2C). PT with TFCE had significantly higher replicability than 3 kinds of cluster-based correction (i.e., GRF, AFNI 3dClustSim and DPABI AlphaSim) with strictest threshold (voxel-wise *P* < 0.0005 and cluster-wise *P* < 0.025, each tail), PT (voxel-wise threshold of *P* < 0.002 (Z > 3.09) with cluster-wise thresholds of *P* < 0.05 (two tailed)), PT with voxel-wise correction (VOX) and FDR in the post-hoc analysis (*P* < 0.05, multiple comparison corrected by Tukey's honest significant difference criterion) (Figure 2C).

**Table 5:**
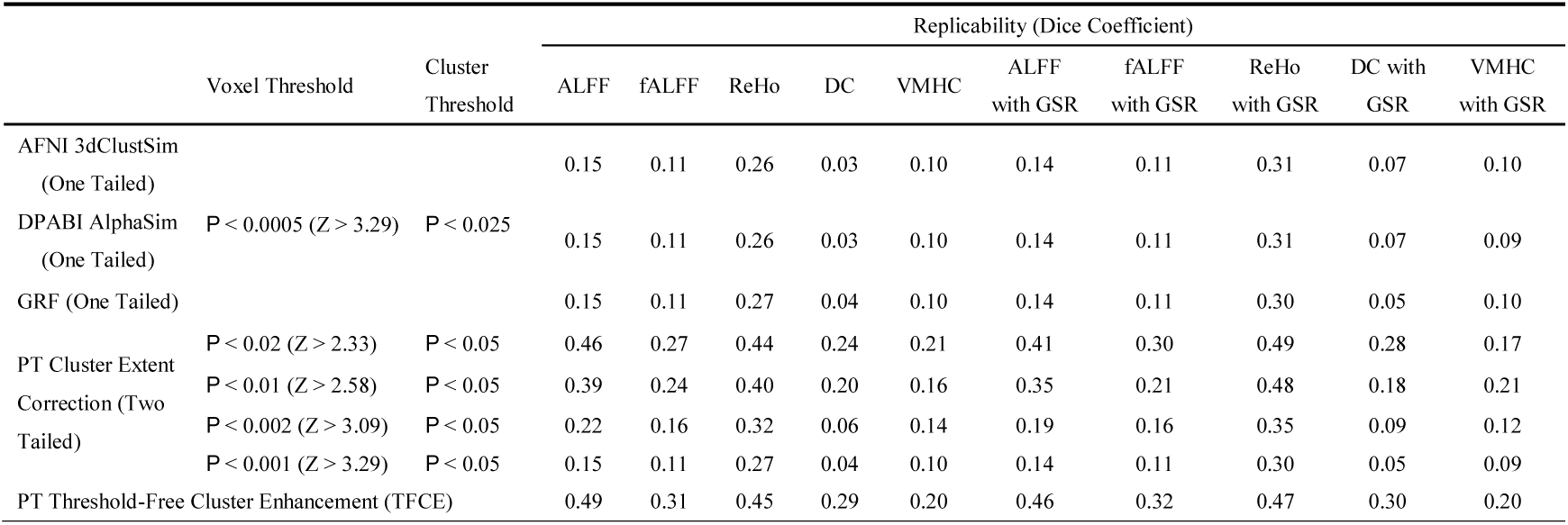
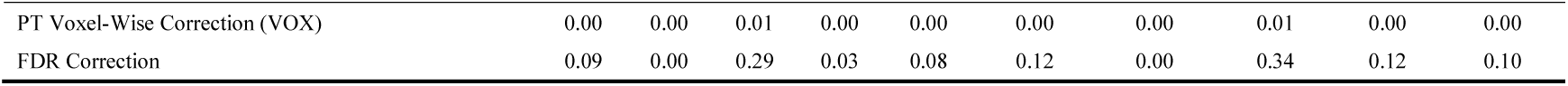
Replicability of eyes-open eyes-closed (EOEC) differences for all R-fMRI metrics with and without Global Signal Regression (GSR) under correction of 3 kinds of cluster-based correction with the strictest threshold, 6 kinds of Permutation Test (PT) based correction and False Discovery Rate (FDR) correction, calculated using significant results in Beijing EOEC1 and EOEC2 datasets. For replicability for all the 31 kinds of multiple comparison correction strategies, please see Table S15.

|  |  |  | Replicability (Dice Coefficient) |  |  |  |  |  |  |  |  |  |
| --- | --- | --- | --- | --- | --- | --- | --- | --- | --- | --- | --- | --- |
|  | Voxel Threshold | Cluster Threshold | ALFF | fALFF | ReHo | DC | VMHC | ALFF with GSR | fALFF with GSR | ReHo with GSR | DC with GSR | VMHC with GSR |
| AFNI 3dClustSim (One Tailed) |  |  | 0.15 | 0.11 | 0.26 | 0.03 | 0.10 | 0.14 | 0.11 | 0.31 | 0.07 | 0.10 |
| DPABI AlphaSim (One Tailed) | $P < 0.0005$ ($Z > 3.29$) | $P < 0.025$ | 0.15 | 0.11 | 0.26 | 0.03 | 0.10 | 0.14 | 0.11 | 0.31 | 0.07 | 0.09 |
| GRF (One Tailed) |  |  | 0.15 | 0.11 | 0.27 | 0.04 | 0.10 | 0.14 | 0.11 | 0.30 | 0.05 | 0.10 |
| PT Cluster Extent | $P < 0.02$ ($Z > 2.33$) | $P < 0.05$ | 0.46 | 0.27 | 0.44 | 0.24 | 0.21 | 0.41 | 0.30 | 0.49 | 0.28 | 0.17 |
| Correction (Two Tailed) | $P < 0.01$ ($Z > 2.58$) | $P < 0.05$ | 0.39 | 0.24 | 0.40 | 0.20 | 0.16 | 0.35 | 0.21 | 0.48 | 0.18 | 0.21 |
| | $P < 0.002$ ($Z > 3.09$) | $P < 0.05$ | 0.22 | 0.16 | 0.32 | 0.06 | 0.14 | 0.19 | 0.16 | 0.35 | 0.09 | 0.12 |
| | $P < 0.001$ ($Z > 3.29$) | $P < 0.05$ | 0.15 | 0.11 | 0.27 | 0.04 | 0.10 | 0.14 | 0.11 | 0.30 | 0.05 | 0.09 |
| PT Threshold-Free Cluster Enhancement (TFCE) |  |  | 0.49 | 0.31 | 0.45 | 0.29 | 0.20 | 0.46 | 0.32 | 0.47 | 0.30 | 0.20 |

|  |  |  |  |  |  |  |  |  |  |  |
| --- | --- | --- | --- | --- | --- | --- | --- | --- | --- | --- |
| PT Voxel-Wise Correction (VOX) | 0.00 | 0.00 | 0.01 | 0.00 | 0.00 | 0.00 | 0.00 | 0.01 | 0.00 | 0.00 |
| FDR Correction | 0.09 | 0.00 | 0.29 | 0.03 | 0.08 | 0.12 | 0.00 | 0.34 | 0.12 | 0.10 |

We further analyzed the spatial locations of significant EOEC differences. As illustrated in Figure 4, replicable significant EOEC differences were observed mainly in bilateral precentral and postcentral gyrus (EC > EO) and bilateral occipital cortices (EO > EC).

**Figure 4.**
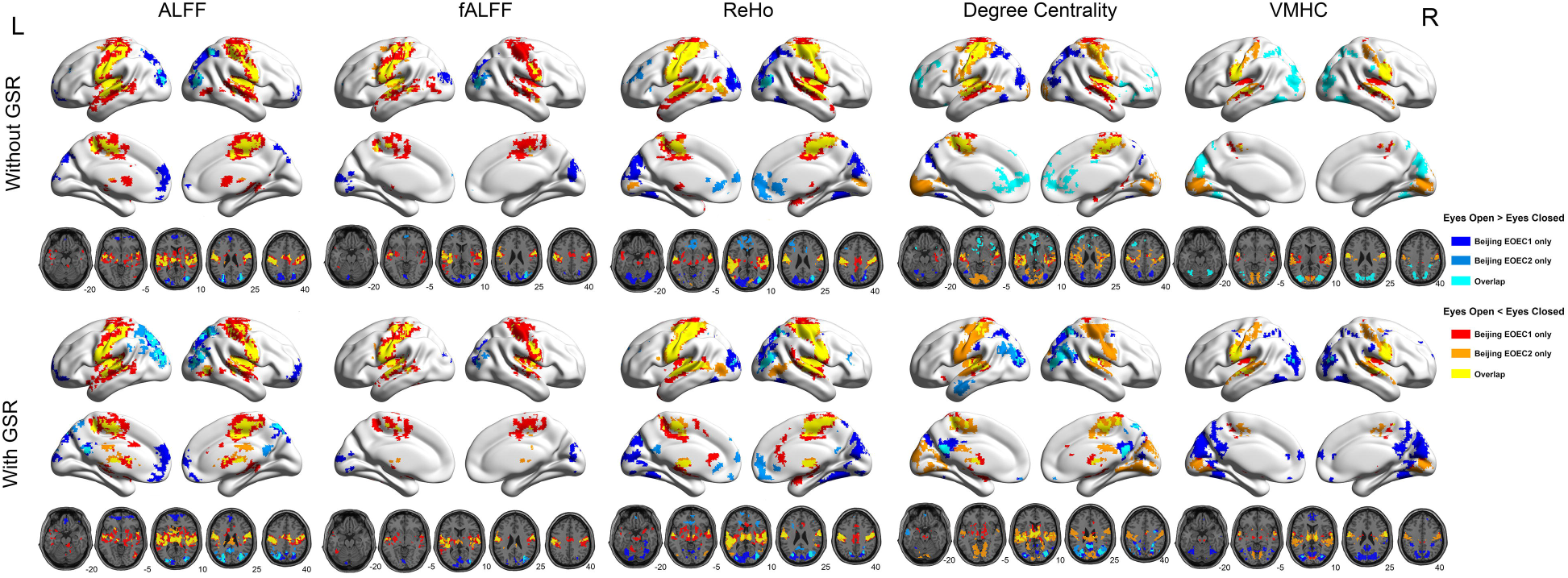
Eyes-open eyes-closed (EOEC) differences those are significant in 2 EOEC datasets, under the correction of Permutation Test (PT) with Threshold-Free Cluster Enhancement (TFCE). Different colors indicate voxels’ EOEC differences are significant in only one dataset (dark color) or in both datasets (bright color).

### 3.6. Influences of Sample Size on Test-retest Reliability, Sensitivity and PPV

First, we assessed the test-retest reliability of sex differences in ALFF without GSR across different sample sizes (k), which we measured using the Dice coefficient (Figure 5A, Table 6). Mean test-retest reliability gradually increased from 0.02 (Dice coefficient, SD = 0.08, k = 30) to 0.46 (Dice coefficient, SD = 0.07, k = 200). However, at a classical sample size for R-fMRI (k = 60, 30 per group), mean test-retest reliability was only 0.08 (Dice coefficient, Table 6).

**Table 6:** Test-retest reliability, sensitivity and positive predictive value (PPV) on ALFF (without GSR) across different sample sizes (k). Both mean and standard deviation (SD) across 100 randomizations were listed.

| Sample Size (k) | Test-retest Reliability<br>(Dice Index) | Sensitivity | PPV |
| --- | --- | --- | --- |
| 30 | 0.02±0.08 | 0.001±0.004 | 0.02±0.09 |
| 40 | 0.03±0.11 | 0.001±0.01 | 0.07±0.21 |
| 50 | 0.05±0.13 | 0.004±0.01 | 0.07±0.19 |
| 60 | 0.08±0.17 | 0.01±0.02 | 0.12±0.22 |
| 70 | 0.16±0.21 | 0.01±0.02 | 0.17±0.22 |
| 80 | 0.23±0.22 | 0.02±0.03 | 0.26±0.24 |
| 90 | 0.28±0.21 | 0.04±0.04 | 0.25±0.16 |
| 100 | 0.32±0.19 | 0.05±0.04 | 0.28±0.14 |
| 120 | 0.36±0.14 | 0.10±0.06 | 0.29±0.08 |
| 140 | 0.39±0.11 | 0.17±0.08 | 0.29±0.04 |
| 160 | 0.39±0.09 | 0.23±0.09 | 0.30±0.03 |
| 180 | 0.42±0.08 | 0.32±0.09 | 0.30±0.02 |
| 200 | 0.46±0.07 | 0.43±0.07 | 0.30±0.02 |

**Figure 5.**
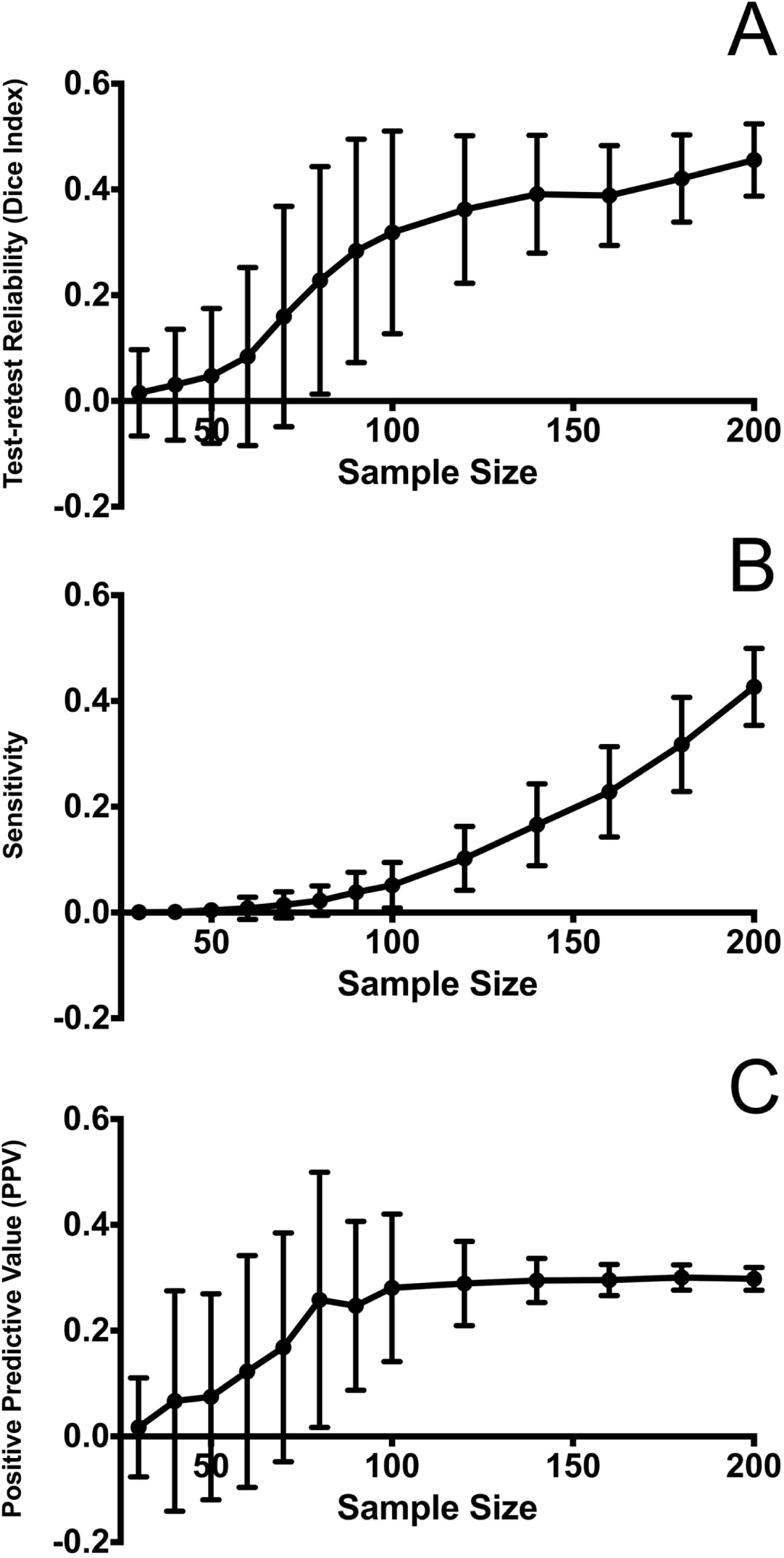
Test-retest reliability (Dice index), sensitivity and positive predictive value (PPV) on ALFF (without GSR) as functions of sample size.

For significant voxels in both tests of each randomization and each k, we calculated sensitivity (power) and PPV using the previously defined “gold standard” (significant voxels in both CORR sessions and in the FCP dataset after correction of PT with TFCE). As shown in Figure 5B and Table 6, mean sensitivity increased from 0.0007 (SD = 0.004, k = 30) to 0.43 (SD = 0.07, k = 200). For PPV, after increasing from 0.02 (SD = 0.09, k = 30) to 0.26 (SD = 0.24, k = 80), PPV reached an asymptote at around 0.30 (Figure 5C, Table 6).

## 4. DISCUSSION

A recent analysis observed that the conclusions drawn from many neuroimaging studies are probably irreproducible (Poldrack et al., 2017). Lack of reproducibility may partly be due to (a) the abuse of liberal multiple comparison correction strategies and (b) the high prevalence of small sample size studies. Here, we provided a comprehensive examination of the impact of different multiple comparison correction strategies and sample size on test-retest reliability and replicability across widely used R-fMRI metrics. We found that multiple comparison correction strategies with liberal thresholds could yield higher test-retest reliability and replicability but at the cost of dramatically increasing FWER to unacceptable levels. We noted that permutation test with TFCE, a strict multiple comparison correction strategy, reached the best balance between FWER (under 5%) and test-retest reliability and replicability (e.g., 0.68 test-retest reliability and 0.25 replicability of sex differences in ALFF without GSR). Although sex differences in R-fMRI metrics could be detected with moderate test-retest reliabilities, they were poorly replicated in a distinct dataset (replicability of sex differences < 0.3). Among the brain regions showing the most reproducible sex differences, PCC demonstrated consistently lower spontaneous activity in males compared with females. Furthermore, by calculating replicabilities with two independent within-subject EOEC datasets, we found that the better performance of permutation test with TFCE generalized to within-subject design studies. Defining the most reproducible brain regions in two large sample datasets as a “gold standard”, and randomly drawing subjects with different sample sizes from one single site, we found that both test-retest reliability and sensitivity increased with sample size. However, PPV reached a plateau at k=80 (40 per group) and remained around 0.30 even with further sample size increases. Here we discuss the implications of our findings on decision-making regarding the choice of multiple comparison correction strategies and approach towards addressing the challenge of reproducibility.

### 4.1. Selecting a Multiple Comparison Correction Strategy with Respect to FWER

Appropriate multiple comparison correction strategies must control the false positive rate at an acceptable level. Our results replicated the findings of prior work (Eklund, et al., 2016), which analyzed R-fMRI data with a putative task design to compute FWER in task fMRI studies. They also performed between-group comparisons on simulated null task activation maps and calculated the FWER. They found an unacceptably high FWER for most widely used multiple comparison correction strategies. Our results provide additional evidence from group comparisons with a range of R-fMRI metrics. Our results confirmed that multiple comparison correction strategies with a liberal threshold (e.g., with voxel wise *P* < 0.01 and cluster wise *P* < 0.05) led to an unacceptably high FWER, while PT can maintain the FWER at nominal 0.05 levels.

Beyond replicating Eklund et al.’s conclusions regarding FWER, two additional points should be noted. First, researchers should pay close attention to whether the test is one-tailed or two-tailed. As most researchers are interested in two-tailed effects (e.g., both patients > controls and patients < controls), if they perform one-tailed thresholding twice (i.e., each tail *P* < 0.05), then the final FWER will be higher than 10% even if the voxel-level p is set to 0.0005 (*Z* > 3.29). Such researchers have to correct for the two tests at each tail, that is, researchers could perform one-tailed correction twice, with each tail voxel-wise *P* < 0.0005 and cluster-wise *P* < 0.025. With such a setting in GRF correction, the FWER almost reaches the nominal level of 5%. Second, among the cluster-based thresholding strategies of GRF, AFNI 3dClustSim and DPABI AlphaSim, we recommend the use of GRF. At the strictest level (voxel-wise *P* < 0.0005 and cluster-wise *P* < 0.025, each tail), GRF is almost valid, while Monte Carlo Simulation based corrections (AFNI 3dClustSim and DPABI AlphaSim) still cannot control FWER at nominal 5% level in some cases. DPABI AlphaSim should not be used in the future, as it always gives looser cluster size threshold and higher FWER, even at a cost of high computational demand. Furthermore, several new options and programs, such as the “ACF” option implemented in 3dClustSim, 3dLocalACF and the “ETAC” option in 3dXClustSim have been proposed to overcome deficits pointed out by Eklund et al. However, according to the recent redux by the AFNI group, these approaches were either inefficient in reducing FWERs or still under development (Cox, et al., 2017). Thus we did not apply these new methods in the current study, but we believe these efforts deserve further investigation in future work.

In sum, in considering FWER, eight different multiple comparison correction strategies can be used: 1) GRF correction with strict p values (voxel wise *P* < 0.0005 and cluster wise *P* < 0.025 for each tail); 2) four kinds of PT with extent thresholding; 3) PT with TFCE; 4) PT with voxel-wise correction; and 5) FDR correction.

### 4.2. Selecting a Multiple Comparison Correction Strategy with Regard to Test-retest Reliability and Replicability

FWER is not the only criterion in selecting a multiple comparison correction strategy; test-retest reliability and replicability may be even more crucial. An appropriate strategy should best balance FWER and reproducibility. For example, GRF with liberal thresholds (e.g., with voxel wise *P* < 0.01 and cluster wise *P* < 0.05) has relatively high test-retest reliability and replicability, but it has unacceptably high FWER. On the other hand, PT with voxel-wise correction can control FWER at a low level (< 5%), but results in the lowest test-retest reliability and replicability, thus it is also not an appropriate strategy to correct for multiple comparisons. Fortunately, PT with TFCE provides a good balance between FWER and reproducibility. PT with TFCE can maintain the FWER under 5%, while yielding moderate test-retest reliability and replicability, e.g., 0.68 test-retest reliability for ALFF on sex differences. Of note, test-retest reliability (sex differences) and replicability (both sex differences and EOEC differences) of PT with TFCE were not significantly lower than for the liberal GRF threshold (e.g., with voxel wise *P* < 0.01 and cluster wise *P* < 0.05).

In considering both FWER as well as test-retest reliability and replicability in both between-subject and within-subject design studies, we recommend using PT with TFCE. As an approach for defining a cluster-like voxel-wise statistic, TFCE avoids the limitation of defining the initial cluster-forming threshold as do other common cluster-based strategy thresholding strategies (Smith and Nichols, 2009). TFCE uses the height parameter (H) and the extent parameter (E) to enhance cluster-like features in a statistical image. Although tweaking of these two parameters is possible, we found the default parameters (H = 2, E = 0.5) already perform well. Of note, PT with TFCE can be easily performed for many different kinds of statistical tests in DPABI, which integrated functions from PALM (Winkler, et al., 2016).

### 4.3. Are R-fMRI Findings Reproducible?

Concerns regarding the reproducibility of R-fMRI findings are increasing (Poldrack et al., 2017). Assessing reproducibility is highly sensitive to the statistical threshold used to define significance (Rombouts, et al., 1998). After identifying the appropriate statistical approach (PT with TFCE), we could evaluate two important aspects of reproducibility, test-retest reliability and replicability, in common R-fMRI metrics. We found most R-fMRI metrics demonstrated moderate test-retest reliability in between-subject contrasts of sex differences (Table 3). Specifically, when computed without GSR, fALFF reached the highest test-retest reliability (0.75), followed by ALFF (0.68) and ReHo (0.54), under the correction of PT with TFCE. DC (0.48) and VMHC (0.44) had the lowest test-retest reliabilities in between-subject comparisons of sex differences. This is consistent with prior studies which reported test-retest reliabilities of R-fMRI networks localized by either seed based analysis (Kristo, et al., 2014) or independent component analysis (Meindl, et al., 2010; Pinter, et al., 2016), showing moderate to high test-retest reliability (between 0.29 and 0.76 in most regions).

Beyond test-retest reliability, a unique contribution of our study is the investigation of replicability. That is, to what extent can a finding in one dataset (usually one study) be replicated in another dataset (another study)? We found the between-subject replicability of sex differences was much lower than test-retest reliability: replicability of all the R-fMRI metrics was below 0.3. ALFF had the best balance between test-retest reliability (0.68) and replicability (0.25), outperforming the other R-fMRI metrics. Although fALFF reached a high test-retest reliability, its replicability was poor (0.06), possibly because it is sensitive to variations in repetition time (TR) used in different datasets. Such low replicability should not be surprising, given the substantial differences between two different datasets, e.g., variation in ethnicity, sequence type, coil type, scanning parameters, participant instructions and head-motion restraint techniques. In evaluating replicabilities of within-subject EOEC contrasts, we found better replicability than for between-subject sex differences, likely reflecting the much larger effect sizes of within-subject EOEC differences than between-subject sex differences (Figure 6). However, this was still not adequate as all replicabilities were below 0.5. The best replicability that could be achieved was 0.49 for EOEC differences of ALFF under PT correction with TFCE. The present results question the generalizability of both between-subject and within-subject results reported in R-fMRI studies, and support the suggestion that future studies incorporate advanced data standardization techniques (Yan, et al., 2013b) to improve replicability.

**Figure 6.**
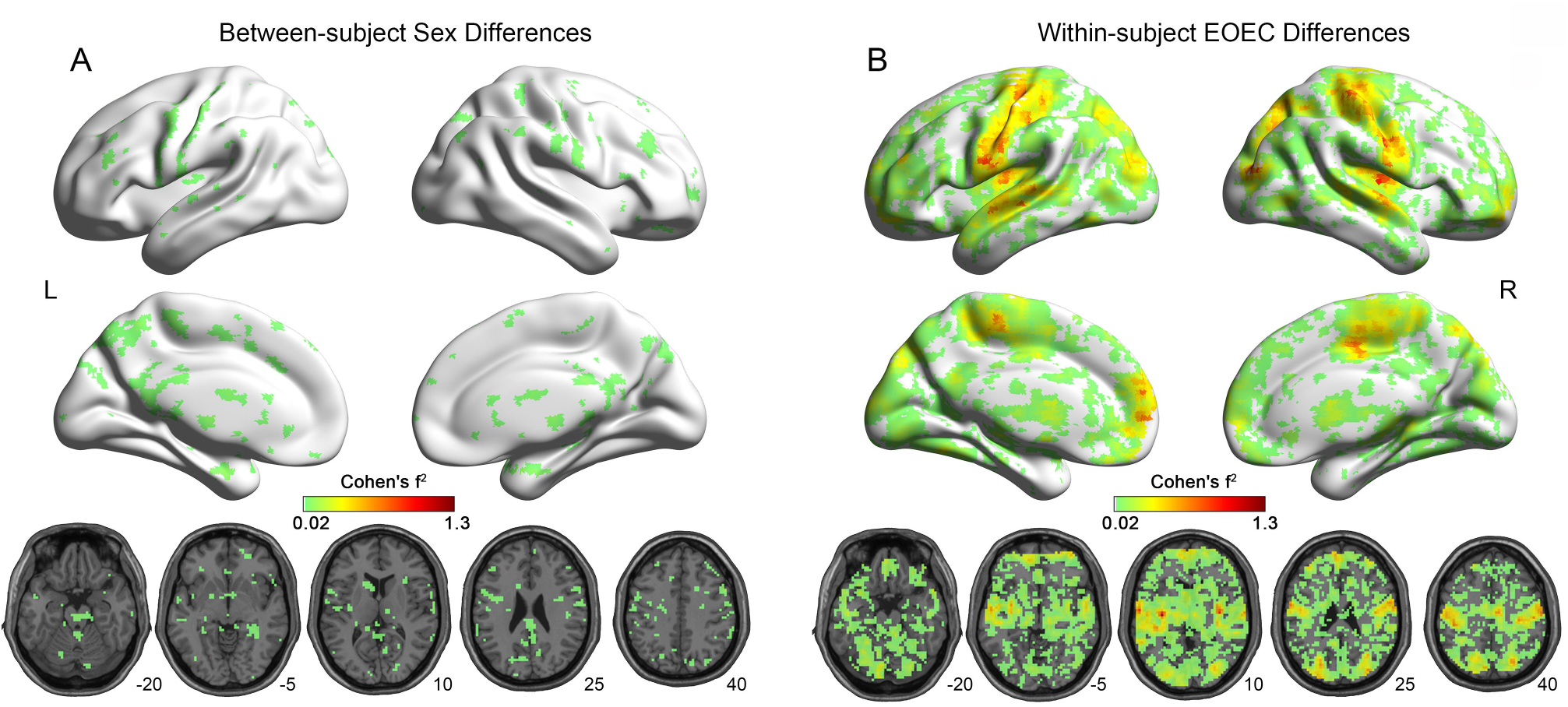
Effect sizes (Cohen’s f^2^) of between-subject sex differences (A: calculated with the first session from CORR dataset, n = 420) and within-subject EOEC differences (B: calculated with the Beijing EOEC1 dataset, n = 48). Cohen’s f^2^ were thresholded at f^2^ > 0.02 (small effect size).

It is noteworthy that we found convergent sex differences and EOEC differences across all metrics and all datasets, despite low replicability. The most replicable sex difference was located in PCC. Greater activity in females versus males was found in PCC, which is similar to previous studies (Allen, et al., 2011; Biswal, et al., 2010). As this phenomenon replicated in two sessions of the same dataset, and was reproduced in two different datasets, we believe this reflects a true sex difference that should be reproducible in future studies. PCC has been shown to be more active in females than in males in several fMRI activation experiments of working and episodic memory (Filippi, et al., 2013). It has been suggested that the PCC is associated with self-referential thoughts, emotions relating to others, remembering the past and thinking about the future (Fransson and Marrelec, 2008; Leech and Sharp, 2014; Maddock, et al., 2001), thus our results are consistent with more inward thinking and empathy in women compared to men. As for EOEC differences, we found greater activity in EC versus EO in precentral and postcentral gyrus and weaker activity in bilateral occipital cortex. Our results were in line with previous studies (Marx, et al., 2004; Yang, et al., 2007), indicating a subtle and important difference in brain activities between these two states.

### 4.4. What Can Be Done for Small Sample Size R-fMRI Studies?

A recent theoretical analysis (Button et al., 2013) highlighted the detrimental effect of low statistical power induced by small sample size on reproducibility. We confirmed that the reliability of results from small sample size studies was very low. For example, under PT with TFCE correction, test-retest reliability was only 0.08 ± 0.17 when k=60 (30 subjects per group), which is a “classical” sample size in the R-fMRI field. According to the mathematical model of bias in scientific research (Button, et al., 2013), studies with a small sample size not only have a reduced chance to detect true effects, but they also reduce the likelihood that a statistically significant result reflects a true effect. The current study used empirical data (R-fMRI metrics) to confirm that the power (sensitivity) of small sample size comparisons is extremely low (around 0.01 when k=60), which is consistent with the finding that the median statistical power across 461 neuroimaging studies was 8% (Button, et al., 2013). Further, small, underpowered samples are more likely to provide positive results through selective analysis and outcome reporting, which are prevalent in R-fMRI studies across a broad range of experimental design and data analytic strategies (Carp, 2012a; Poldrack, et al., 2017). Thus, our results add to the growing consensus in the field calling for larger sample sizes. Indeed, as sample size increased from k=30 to 200, we found reliability increased steadily from 0.02 ± 0.08 to 0.46 ± 0.07, and sensitivity increased from 0.0007 ± 0.0004 to 0.43 ± 0.07. Although PPV reached a plateau at k = 80, it increased from 0.02 ± 0.09 (k=30) to 0.30 ± 0.02 (k=200). Our results quantify the insufficiency of the present classical sample size in the R-fMRI field. For studies examining effect sizes similar to or even less than those of sex differences, results from a sample size less than 80 (40 per group) should be considered preliminary, given their low reliability (< 0.23), sensitivity (< 0.02) and PPV (< 0.26). Alternatively, researchers could prefer within-subject design to between-subject design if it’s possible, as larger effect size in within-subject design can increase reproducibility in small sample size studies, as we demonstrated here in the EOEC difference results.

Another consideration is smoothing kernel in preprocessing. Large smoothing kernels may systematically bias or even obscure evidence of underlying difference at the cost of anatomical specificity (Friston, et al., 1994; Sacchet and Knutson, 2013), thus we chose a relatively small smoothing kernel (4mm FWMH) for preprocessing in the main analyses. We have re-analyzed our data with 8mm FWHM smoothing kernel in preprocessing, which verified our main conclusions in FWER (Figure S1 and Table S12) and the superiority of PT with TFCE (Figure S2 and Tables S16∼S18). Interestingly, we found higher smoothness (8mm in preprocessing) increased test-retest reliability and replicability. For example, for ALFF without GSR under PT with TFCE, replicability of EOEC difference increased from 0.49 to 0.58 (Table 7). Regarding to sample size effect, PPV kept increasing to 0.4 until k = 180 (Figure S3 and Table S19). This indicates larger smoothing kernel (8mm) improves reproducibility and is more likely reflecting true effect. However, finding the optimal smoothing kernel to balance anatomical specificity and reproducibility is out of the scope of the current study, thus needs to be addressed in the future.

**Table 7.** Test-retest reliability and replicability of sex differences, as well as replicability of eyes-open eyes-closed (EOEC) differences under correction of Permutation Test (PT) with Threshold-Free Cluster Enhancement (TFCE), when smoothing kernel in preprocessing was set to 8mm FWMH.

|  | ALFF | fALFF | ReHo | DC | VMHC | ALFF<br>with GSR | fALFF<br>with GSR | ReHo<br>with GSR | DC with<br>GSR | VMHC<br>with GSR |
| --- | --- | --- | --- | --- | --- | --- | --- | --- | --- | --- |
| Test-retest Reliability (Sex difference) | 0.70 | 0.83 | 0.63 | 0.54 | 0.40 | 0.69 | 0.82 | 0.48 | 0.27 | 0.46 |
| Replicability (Sex difference) | 0.35 | 0.50 | 0.22 | 0.20 | 0.00 | 0.33 | 0.54 | 0.16 | 0.05 | 0.00 |
| Replicability (EOEC difference) | 0.58 | 0.46 | 0.45 | 0.33 | 0.17 | 0.55 | 0.49 | 0.53 | 0.32 | 0.18 |

Many suggestions have been proposed to address the challenges of reproducibility, e.g., establishing large-scale consortia to accumulate big data, sharing custom analysis code, following accepted standards for reporting methods, and encouraging replication studies (Button, et al., 2013; Poldrack, et al., 2017). Recently, data-sharing initiatives (e.g., grassroots efforts such as FCP/INDI, openfMRI, fMRIDC and coordinated efforts such as ADNI, HCP, PING and UKBiobank) enable big data research models to address the reproducibility challenge. However, raw data sharing requires intensive coordinating efforts, huge manpower demands and large-capacity data storing/management facilities. Furthermore, sharing raw data entails privacy concerns arising from the possibility of being able to identify participants from high dimensional raw data. These concerns, together with the demands of data organization and the limitation of large data uploading, prevents the wider imaging community from publically sharing valuable brain imaging datasets. The R-fMRI Maps project (http://rfmri.org/maps) was proposed to address the above concerns by only sharing the final maps of various R-fMRI indices, which only need light data storing/uploading requirements and remove the privacy concerns regarding raw data sharing. All of the R-fMRI metric maps of the current study have been made available through the R-fMRI Maps project, thus readers can easily confirm/reanalyze this data with our shared analysis code (https://github.com/Chaogan-Yan/PaperScripts/tree/master/Chen_2017_HBM).

Different from efforts to collect and share unthresholded statistical maps of the human brain (e.g. NeuroVault.org, Gorgolewski, et al., 2015), the metric maps available through the R-fMRI Maps project can be statistically reanalyzed or fed into machine learning algorithms. Through the R-fMRI Maps project, we hope to build an unprecedented big data repository of brain imaging analyses across a wide variety of individuals: including different neurological and psychiatric diseases and disorders, as well as healthy people with different traits. We hope the availability of such a big data repository will help to address the challenge of reproducibility.

## CONCLUSIONS

To our knowledge, this was the first effort to comprehensively evaluate the impact of different strategies to correct for multiple comparisons as well as of sample size on the reproducibility of group differences in R-fMRI metrics. Our results revealed that PT with TFCE, a strict multiple comparison correction strategy, reached the best balance between FWER and test-retest reliability / replicability. We found moderate test-retest reliability of the R-fMRI metrics we assessed. By contrast, replicability was low, bringing into question the generalizability of results reported in R-fMRI studies. Finally, the present research demonstrated that reliability, sensitivity and PPV increase steadily as sample sizes grow. Of note, findings from R-fMRI studies with small sample sizes are poorly reliable, as well as yielding low sensitivity and PPV, which reinforces calls for increasing sample size in future R-fMRI studies.

## ACKNOWLEDGEMENTS

The authors appreciate the editorial assistance and support of Dr. F. Xavier Castellanos. This work was supported by the National Key R&D Program of China (2017YFC1309902), the National Natural Science Foundation of China (81671774 and 81630031), the Hundred Talents Program of the Chinese Academy of Sciences, and Beijing Municipal Science & Technology Commission (Z161100000216152).

## CONFLICT OF INTEREST

The authors declare no competing financial interests.

---

**Figure S1.** Family wise error rates (when smoothing kernel in preprocessing was set to 8mm FWMH) of ALFF (without GSR) under 31 kinds of different multiple comparison correction strategies. AFNI 3dClustSim and DPABI AlphaSim are two versions of Monte Carlo simulation based correction implemented in AFNI and DPABI, separately. GRF, PT and FDR are Gaussian Random Field correction, Permutation Test and False Discovery Rate correction implemented in DPABI, separately. TFCE stands for Threshold-Free Cluster Enhancement and VOX stands for Voxel-Wise Correction. Both of them are correction approaches accompanied with PT. The red solid line shows the nominal 5% positive false positive rate, and the gray dashed line shows its approximate theoretical 95% confidence interval, 3.65%∼6.35%.

**Figure S2.** Results (when smoothing kernel in preprocessing was set to 8mm FWMH) of the Friedman Test of both test-retest reliabilities and replicabilities regarding between-subject sex differences and within-subject eyes-open eyes-closed (EOEC) differences on 5 metrics by 2 preprocessing strategies (with and without GSR) among 3 kinds of cluster-based correction with the strictest threshold, 6 kinds of Permutation Test (PT) based correction and False Discovery Rate (FDR) correction (A: test-retest reliability regarding between-subject sex differences B: replicability regarding between-subject sex differences C: replicability regarding within-subject EOEC differences). Larger median rank numbers represent the better reproducibility compared with other statistical threshold approaches. PT with TFCE is outlined with red, and those are significantly different from PT with TFCE in reproducibility are outlined with yellow (multiple comparison corrected by Tukey's honest significant difference criterion). GRF, PT and FDR stand for Gaussian Random Field correction, Permutation Test and False Discovery Rate correction, separately. All versions of cluster-based corrections are one-tailed *P* values while all versions of PT are two tailed *P* values.

**Figure S3.** Test-retest reliability (Dice index), sensitivity and positive predictive value (PPV) on ALFF (without GSR) as functions of sample size, when smoothing kernel in preprocessing was set to 8mm FWMH.

